# Recovery of dynamics and function in spiking neural networks with closed-loop control

**DOI:** 10.1101/030189

**Authors:** I. Vlachos, T. Deniz, A. Aertsen, A. Kumar

**Affiliations:** Bernstein Center Freiburg, Faculty of Biology, University of Freiburg, Germany; Computational Biology, School of Computer Science and Communication, KTH Royal Institute of Technology, Stockholm, Sweden

## Abstract

There is a growing interest in developing novel brain stimulation methods to control disease-related aberrant neural activity and to address basic neuroscience questions. Conventional methods for manipulating brain activity rely on open-loop approaches that usually lead to excessive stimulation and, crucially, do not restore the original computations performed by the network. Thus, they are often accompanied by undesired side-effects. Here, we introduce delayed feedback control (DFC), a conceptually simple but effective method, to control pathological oscillations in spiking neural networks. Using mathematical analysis and numerical simulations we show that DFC can restore a wide range of aberrant network dynamics either by suppressing or enhancing synchronous irregular activity. Importantly, DFC besides steering the system back to a healthy state, it also recovers the computations performed by the underlying network. Finally, using our theory we isolate the role of single neuron and synapse properties in determining the stability of the closed-loop system.

---

In the past decades open-loop brain stimulation has been used as a common tool to restore aberrant neuronal activity. The most successful example is the application of high-frequency deep-brain-stimulation (DBS) used to ameliorate motor symptoms in Parkinson’s disease (PD). However, even in this case the stimulation induces side-effects such as gait imbalance, cognitive impairment, speech impairment, depression etc^1^. The main cause of these side-effects is likely to be the constant stimulation, but additional explanations are plausible, e.g. the inability of open-loop stimulation to recover the original computations carried out by the impaired brain area. Thus, there is a clear need for more sophisticated brain stimulation schemes^2-4^.

Moreover, to exploit the full potential of external brain stimulation as a research and therapeutic tool it is important to obtain theoretical insights that can guide the design of novel stimulation protocols. The goal for these new stimulation methods should ideally be twofold: to alter the dynamical state of the brain activity in a desired manner and to recover the computations performed by the network. Here, we demonstrate that DFC, an effective feedback method with origins in chaos control^5,6^, achieves this objective.

To show that DFC is effective in altering the global activity state, we focus on its ability to switch the network state between synchronous-irregular (SI) oscillatory and asynchronous-irregular (AI) non-oscillatory activity. This choice is motivated by the fact that several brain diseases are manifested as a transformation of the AI state to persistent SI oscillations, such as in PD^7^ and in certain forms of epilepsy^8^, or as the inability of the network to generate transient SI activity, e.g. in schizophrenia^9^. To demonstrate that DFC facilitates the recovery of certain types of computations, we illustrate how a network under DFC can effectively process and route rate as well as temporally coded signals. Therefore, DFC not only steers the system to a more physiological activity regime, but it also recovers the coding abilities of the network as they were present before the onset of the pathology.

Previous theoretical models of closed-loop stimulation are not suitable to study the control of SI oscillations because the dynamics that arise in networks of phase oscillators^10-12^, in networks of Hodgkin-Huxley neurons^13,14^ and in Wilson-Cowan type of firing rates models^15^ are qualitatively different from the SI oscillations^16,17^. In addition, the physiologically plausible SI oscillations are known to be robust to both noise and heterogeneities^18-20^ and, therefore, require a more differentiated control approach. Finally, the theoretical insights we provide into the mechanisms of feedback control in SNNs could provide an explanation for the recent success of event-driven stimulation schemes^21-23^ as well.

## Results

Excitation and inhibition (EI) in balanced random SNNs cause asynchronous, irregular (AI) and non-oscillatory population activity. This state resembles the ongoing activity in the healthy state^17^. Changes in the EI balance, caused by altered inputs and/or changes in the recurrent synaptic strengths, can result in two qualitatively different types of oscillations. The synchronous-regular (SR) oscillations arise when the mean input to the individual neurons exceeds their spiking threshold, resulting in high firing rate and high frequency regular oscillations^16,17^. By contrast, the synchronous-irregular (SI) oscillations arise because of synaptic strong coupling and increased variance of the total input to the neurons. In the SI state individual neurons spike irregularly at a lower rate than the oscillation frequency. Importantly, the emergence of the SI oscillations is accompanied with a change in the network transfer function and its ability to represent stimulus-related activity. Persistent SI oscillations often are signature of brain diseases, e.g. in PD^7^ and epilepsy^8^. The altered network transfer function and the robustness of the oscillations to noise and neuronal heterogeneities pose a serious challenge for stimulation-based therapeutic approaches. In the following we show that DFC is able to both quench SI oscillations and to recover the original network transfer function.

## Control of SI activity in I-I networks

While our goal is to reveal the mechanisms by which DFC controls SI activity in excitatory-inhibitory SNNs, it is more instructive to first demonstrate the concept in a simple, purely inhibitory SNN. In such a network, the emergence of SI oscillations can be investigated by analyzing the stability of the network firing rate in the AI state^18,19^. A small perturbation of the steady-state firing rate

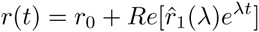

leads to a perturbation in the recurrent input

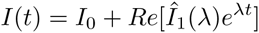

with 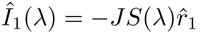, where *J* is the synaptic coupling strength and *S* is the synaptic response function. Both perturbations have to be consistent, that is

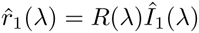

where *R* is the neuron response function. This results in:

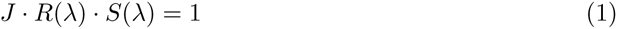

In a purely inhibitory network *J* is negative, but here the negative sign of *J* has been absorbed in the phase *S*(*λ*). We can then compute the eigenvalue spectrum, that is the roots A that satisfy (1). When the eigenvalues have a positive real part, the AI state is unstable and the SNN settles in the SI state. Note that due to the synaptic delays the spectrum is infinite. However, in time-delay systems of the retarded type that we are considering here, the total number of unstable eigenvalues is always finite^24^. Increasing *J* shifts the spectrum towards more positive values on the real axis. For a critical value *J_cr_* a complex pair of eigenvalues crosses the imaginary axis and the system becomes unstable through a supercritical Hopf bifurcation^18^ (**Figure 1**). In the following we consider an SNN in which *J* > *J_cr_* resulting in SI oscillations. We aim at designing a controller that can alter the global activity state from SI to AI by placing the unstable eigenvalues back to the left half-plane (**Figure 1A**).

**Figure 1.**
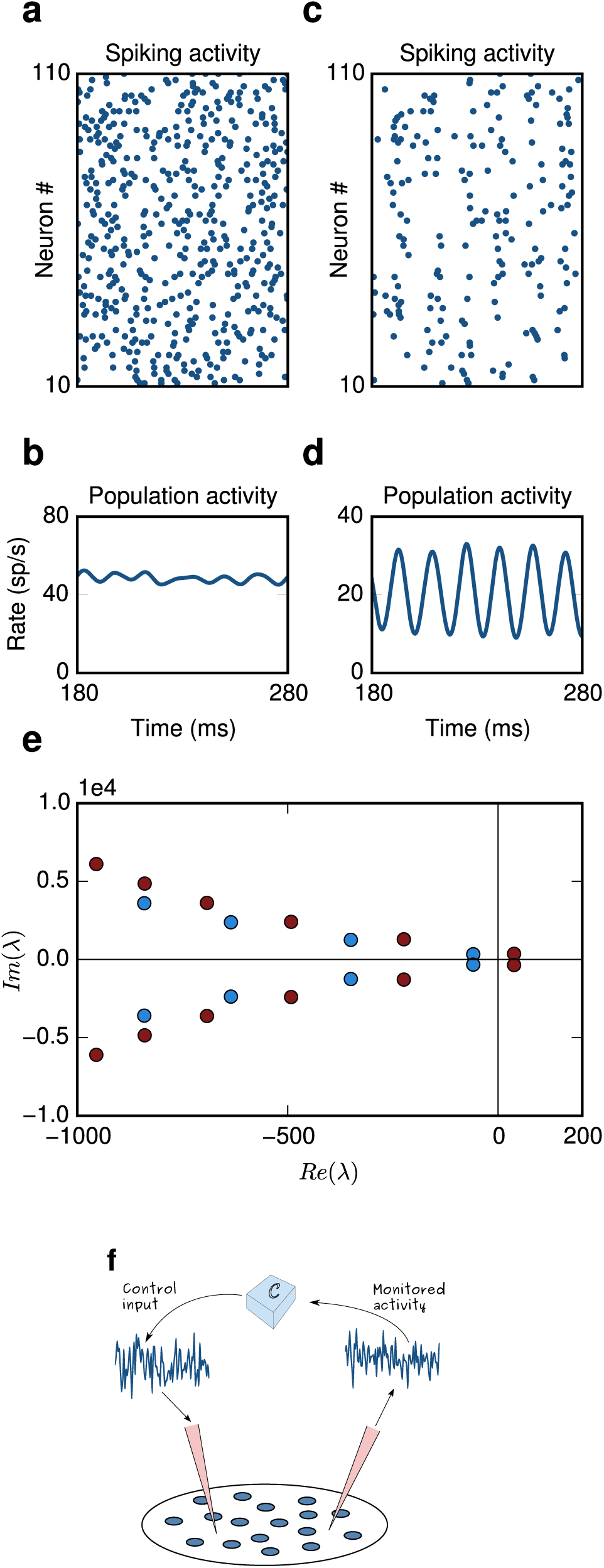
Generation of stochastic oscillations. (**A**) The network is in an asynchronous iregular (AI) regime. Single neuron firing follows Poisson statistics and (**B**) the population activity is stationary. (**C**) The network generates stochastic oscillations. Single neuron firing is still iregular, (**D**) but the population activity is oscillatory. (**E**) Eigenvalue spectrum computed from equation 1. The emergence of oscillations can be explained by the onset of a Hopf bifurcation. When a complex pair of eigenvalues crosses the imaginary axis the network activity becomes unstable. (blue dots: AI regime, red dots: SI oscillations) (**F**) Schematic illustrating closed loop control. The neural activity is being continuously monitored and processed by the controller C, which dynamically generates an appropriate signal that is fed back to the network.

We want to stimulate the network in a closed-loop to alter the SI activity, thus we need to modify equation (1) to include the contribution due to DFC:

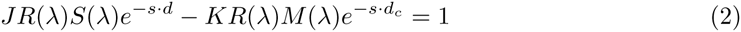

where *K* is the control gain, *d_c_* the control delay and *M* the control kernel. The roots of the above equation (see methods) yield the range of parameters *K, d_c_* that move the unstable eigenvalues back to the left-half plane (**Figure 1A**), which results in a switch of activity from SI to AI.

We simulated a population of 10,000 sparsely connected leaky-integrate-and-fire (LIF) neurons coupled with inhibitory synapses. Consistent with the analytical predictions from the mean-field approximation, for a critical coupling value *J_cr_* the asynchronous irregular (AI) activity destabilizes and stochastic oscillations emerge. Switching on the DFC with parameters estimated from equation (2), almost immediately results in suppression of oscillations and in a network state that resembles the AI regime (**Figure 2A-D**). The suppression of stochastic oscillations is evident both in the spiking activity of single neurons (**Figure 2A**) and in the network population activity (**Figure 2B**). The spike count variability and the irregularity of single neuron interspike intervals, estimated by the Fano Factor (FF) and the coefficient of variation (CV) respectively, confirm that under DFC the firing of individual neurons in the network follows Poisson statistics (AI: *FF* = 1.04, *CV* = 1.01, DFC: *FF* = 1.02, *CV* = 0.99). Moreover, the oscillation index that captures the degree of oscillatory activity (see methods) is in both conditions comparable (AI:*P_T_* = 1.47, DFC: *P_T_* = 1.45) and significantly smaller than in the SI state (*P_T_* = 3). The change in the network spiking activity is also observed in the subthreshold membrane potential of individual neurons (**Figure 2C,D**).

**Figure 2.**
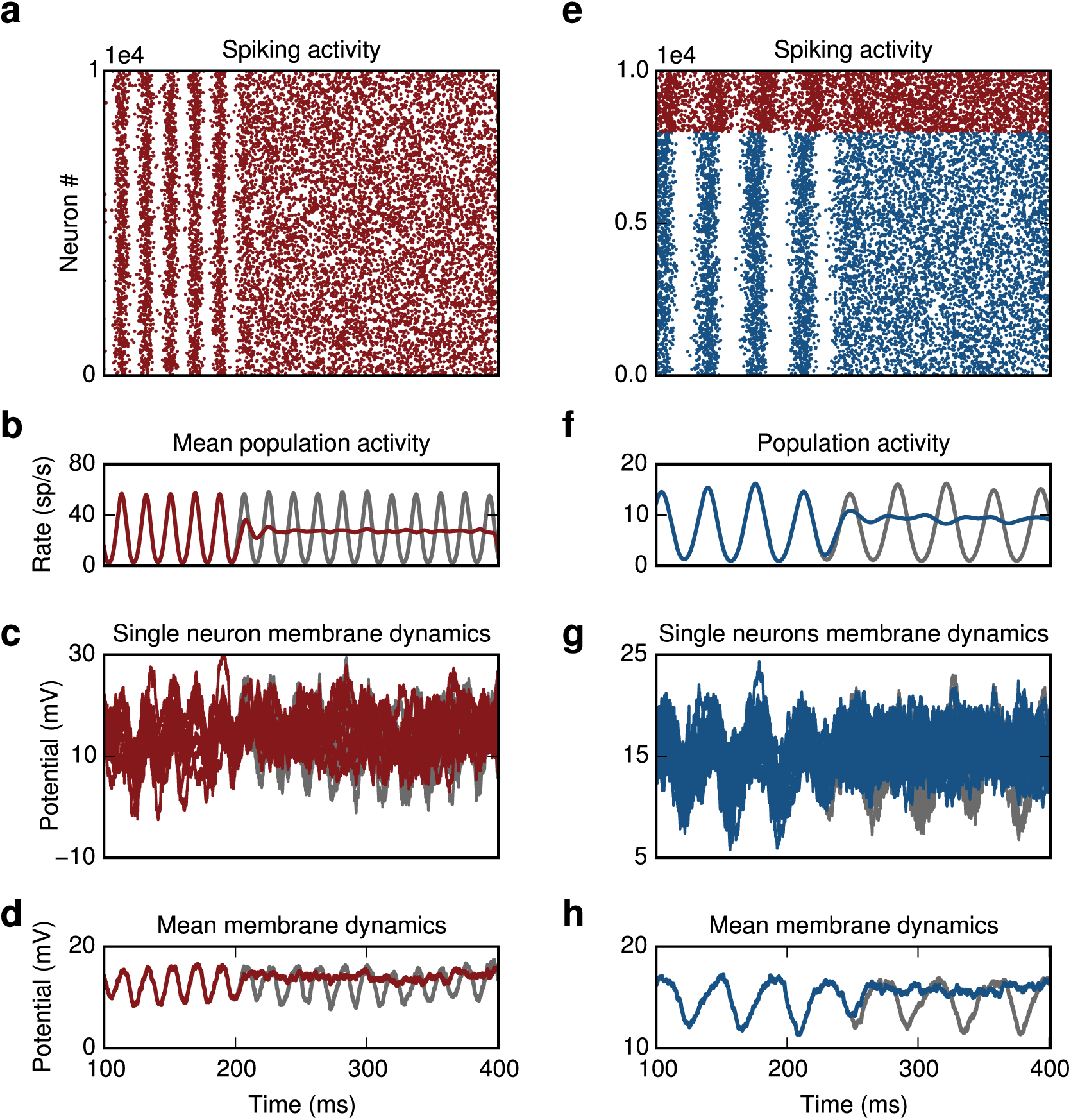
Closed-loop control of oscillations. (**A**) Inhibitory network. Switching on the controller at t=200 ms leads to suppression of oscillations. (**B**) Population activity without (grey) and with control (red). (**C**) Single membrane potential trajectories of ten randomly chosen neurons in the network (**D**) Averaged trace of subthreshold dynamics. (**E**) Excitatory-Inhibitory network. Switching on the controller at t=250 ms leads to suppression of oscillations. (**F**) Activity of excitatory population without (grey) and with control (blue). (**G** and **H**) Same as (**C** and **D**), now membrane potential of excitatory neurons is shown. For better visualization the spike trains in **A** and **E** are thinned out.

## Control of SI activity in E-I networks

Next we demonstrate the applicability of DFC in changing the SI state in recurrent networks of excitatory and inhibitory neurons. To this end we simulated a SNN composed of 8000 excitatory and 2000 inhibitory neurons and tuned the parameters to get an SI state. The self-consistency equation for the coupled EI-network is given by

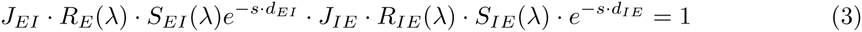

where *J_ij_, d_ij_* is the synaptic coupling strength and delay from population *j* to population *i* and *R_E_* (*R_I_*) is the neuron response function of excitatory (inhibitory) neurons. We implemented DFC by recording the activity of neurons in the inhibitory population while stimulating excitatory neurons. Switching on the controller yielded a near instantaneous transition in the network activity from SI to AI (**Figure 2E-H**). In this case the original physiological state we want to recover was characterized by slightly less irregular firing of the individual neurons. Nevertheless, DFC successfully steered the network to a regime with statistics comparable to the AI activity (DFC: *FF_E_* = 0.85, *CV_E_* = 0.91, AI: *FF_E_* = 0.83, *CV_E_* = 1.03).

In a coupled network with more than one population additional possibilities for recording and stimulating neurons exist. For instance, we could both record and stimulate the excitatory population (see below “stability and robustness of control domains”). Our results, however, do not depend on the exact identity of the recorded and stimulated neurons. The oscillation frequency in the SI state of the network was in the beta range, (*f* ~ 30Hz), which is characteristic for PD^7^. This suggests that if one of the factors contributing to oscillations in PD is strong coupling between STN and GPe^25,26^, then this control approach could be used to suppress these beta band oscillations. The results presented here are general and the same approach can be applied to suppress oscillations in other frequency bands as well.

## Stability and robustness of control domains

To determine the range of values that led to stable control we fixed the control kernel *M*, using a box function of width 1 ms, and parametrized the system by the gain *K* and delay *d_C_.* For each pair of values we simulated the SNN and computed the oscillation index. The (*K, d_c_*)-plane shows that a stable control domain exists at ms. That is, an effective control delay of *d_c_,_eff_* = 7 ms yields the maximum stability for the resulting AI state. The semi-analytical results derived from mean-field theory are in good agreement with the numerical simulations. The only discrepancy occurs when the difference between synaptic and control coupling is small. In this case it is more difficult to maintain constant rates of the stimulated population and the system may become effectively excitatory leading to rate instabilities^27^. Moreover, fluctuations in the mean input that are ignored in our mean-field approach could also become more important. The analysis of these fluctuations is beyond the scope of this work and will be addressed in a future study.

## Differential control

Despite the fact that one stable control domain exists, a compensation mechanism to maintain constant firing rates is required to achieve stable control. In real-life applications a detailed fine-tuning may not always be possible. Therefore, we modified our control protocol and introduced an additional delay term *d_C_*_2_, thus, effectively feeding into the controller the difference between two time-delayed versions of the population activity. For such differential DFC scheme the control signal is given by (see methods):

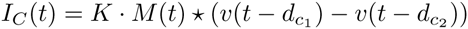

Differential control has been previously used to control unstable periodic orbits^5,28^ and to suppress synchrony in networks with discrete-time neuron models^29^. Here, we accounted for the fact that recording neural activity and injecting a control current into the neurons introduces a finite time-delay. Therefore, we used a small but non-zero value for *d_C_*_2_, i.e. *d_C_*_2_ = 1 ms, which is close to the feedback delays introduced by current technologies^30,31^. A crucial advantage of differential DFC is that no additional rate compensation is required, because the mean contribution of the control signal vanishes

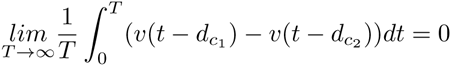

Moving in the control parameter space (*K, d_c_*), therefore, did not affect the firing rates of the neurons. This was reflected in the near perfect overlap of theoretical predictions and numerical simulations of the SNN (**Figure 3B**). In addition, differential DFC introduced two positive effects on the stability of the control domains: (i) The first domain was expanded, which amounts to an increase in the robustness in the parameter variation. That is, small deviations from the estimated values of the gain and the delay would not be critical for the stability of the AI state. (ii) A new stable control domain appeared at *t* =23 ms. Thus, with differential control there is an increase of the range of parameters that lead to stability.

**Figure 3.**
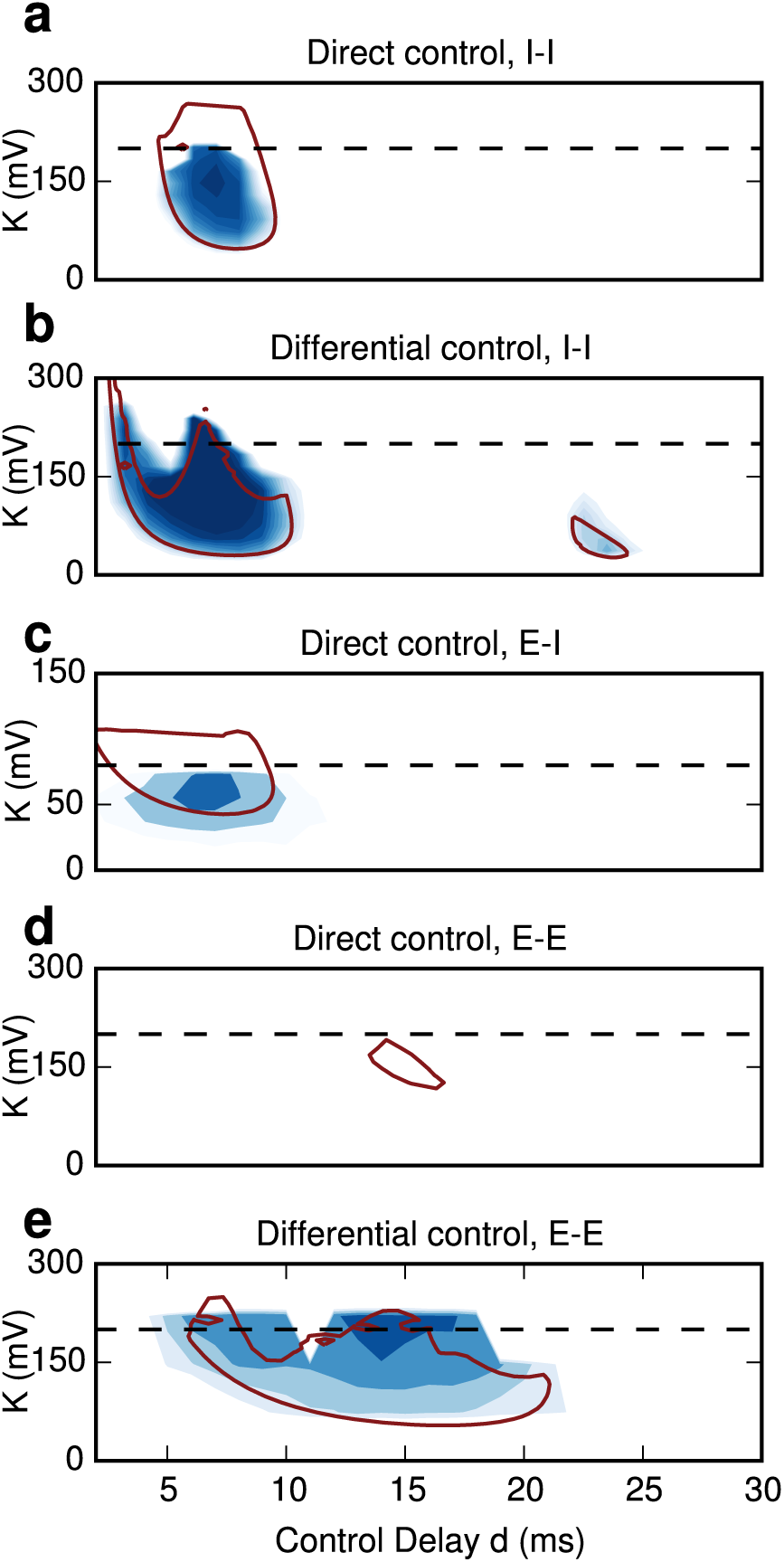
Stability landscape of the network actvitity under DFC. (**A**) Direct DFC in the inhibitory network. One stable control domain appears at t = 7 ms. The results from mean field theory correctly predict the location and shape of the domain (dashed line). The discrepancy for K>200 mV is explained by a deviation of the firing rates between numerical simulations and mean-field theory. (**B**) Differential DFC in the inhibitory network. The first stable control domain around 7 ms is enlarged. An additional small, but stable control domains appears around t=23 ms. Moving in the state space does not affect the firing rates and therefore no rate compensation is required. The numerical and analytical results (red contour) are in perfect agreement. (**C**) Direct DFC in an E-I network. The excitatory population is being stimulated while the activity of the inhibitory neurons is recorded. A stable control domain appears around t=7 ms which reflects the effective delay from the inhibitory to the excitatory population. (**D**) Same as (**C**) but here the excitatory population is both recorded and stimulated at the same time. The theoretical analysis yields a small but stable control domain around t=15 ms. The location of the domains reflects the larger E-I-E loop (see text). In the numerical simulations this control domain does not arise, because fluctuations, which are ignored by the mean field-approach, quickly destabilize the system. (**E**) Same as (**D**) but differential DFC is used. The stable control domain is enlarged showing that differential DFC yields more robust control in coupled E-I networks as well. In all panels regions of stable activity are indicated by shades of blue color. The dashed line denote that total network coupling J.

DFC also enhanced the robustness of the system to external disturbances, e.g. undesired signals at the controller output, measurement noise etc. This becomes evident when we consider the distance *B^cr^* of the complex eigenvalues *λ_i_* from the imaginary axis for the main stable control domain at *t* = 7 ms. The higher the values of *B^cr^* are the more robust the closed-loop system is. Direct and differential control yielded 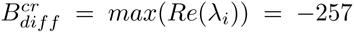 and 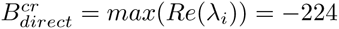 respectively, clearly revealing a more robust system with differential DFC.

Both direct and differential control were effective in coupled E-I populations as well. The location of the stable control domains depended on the exact implementation (**Figure 3**). When the activity of the inhibitory population was monitored while the excitatory population stimulated the main stable control domain appeared at *t* = 7 ms (**Figure 3C**). This location is identical with the purely inhibitory network and reflects the overall delay of the I-E path (I-I loop) in the E-I (I-I network). Indeed for both the I-E path and E-I loop the effective delay is *d_eff_* = 7 ms (see methods). By contrast, when the excitatory population was both recorded and stimulated then the location of the domains shifted to around *t* = 15 ms reflecting the larger overall delay in the E-I-E loop. (**Figure 3D,E**). Note that in this case the stable control domain for direct control was smaller. The reason is that the size of the stable control domains shrinks for larger delays.

## DFC control vs noise injection

In both the I-I and E-I network we applied an identical control signal to all stimulated neurons. That is, we did not disrupt oscillations and decorrelated network activity by injecting different currents to each of the neurons. This is in contrast with a wide-held assumption that common input always tends to increase correlations in neural activity^32^. The results from the application of DFC reveal that common input can both increase or decrease correlations in SNNs. It is the timing and amplitude of the common input that determines the direction in which correlations are affected.

It is important to point out that injection of a control signal is not equivalent to the application of additive noise to the system. To demonstrate this we simulated an I-I network and injected Gaussian noise with the same mean and variance as the control signal to all neurons. This stimulation approach failed to suppress SI oscillations (**Figure 4A,B**) indicating that the temporal structure of the control signal is crucial for successful control. Increasing further the noise intensity, e.g. by a factor of ten, eventually resulted in desynchronization of the activity and in quenching of oscillations (**Figure 4D,E**). However, with such strong strong external noise the network dynamics is predominantly influenced by the input rather than the recurrent activity. This condition is disastrous from a computational point of view, because any information processing taking place within the stimulated brain region would be severely impaired. To illustrate this we recorded the subthreshold dynamics of ten randomly selected neurons in the network (**Figure 4F**). The huge fluctuations in the membrane potential under the influence of strong external noise are rather pathological. By contrast, the fluctuations in the case of DFC are comparable to those in the physiological AI regime.

**Figure 4.**
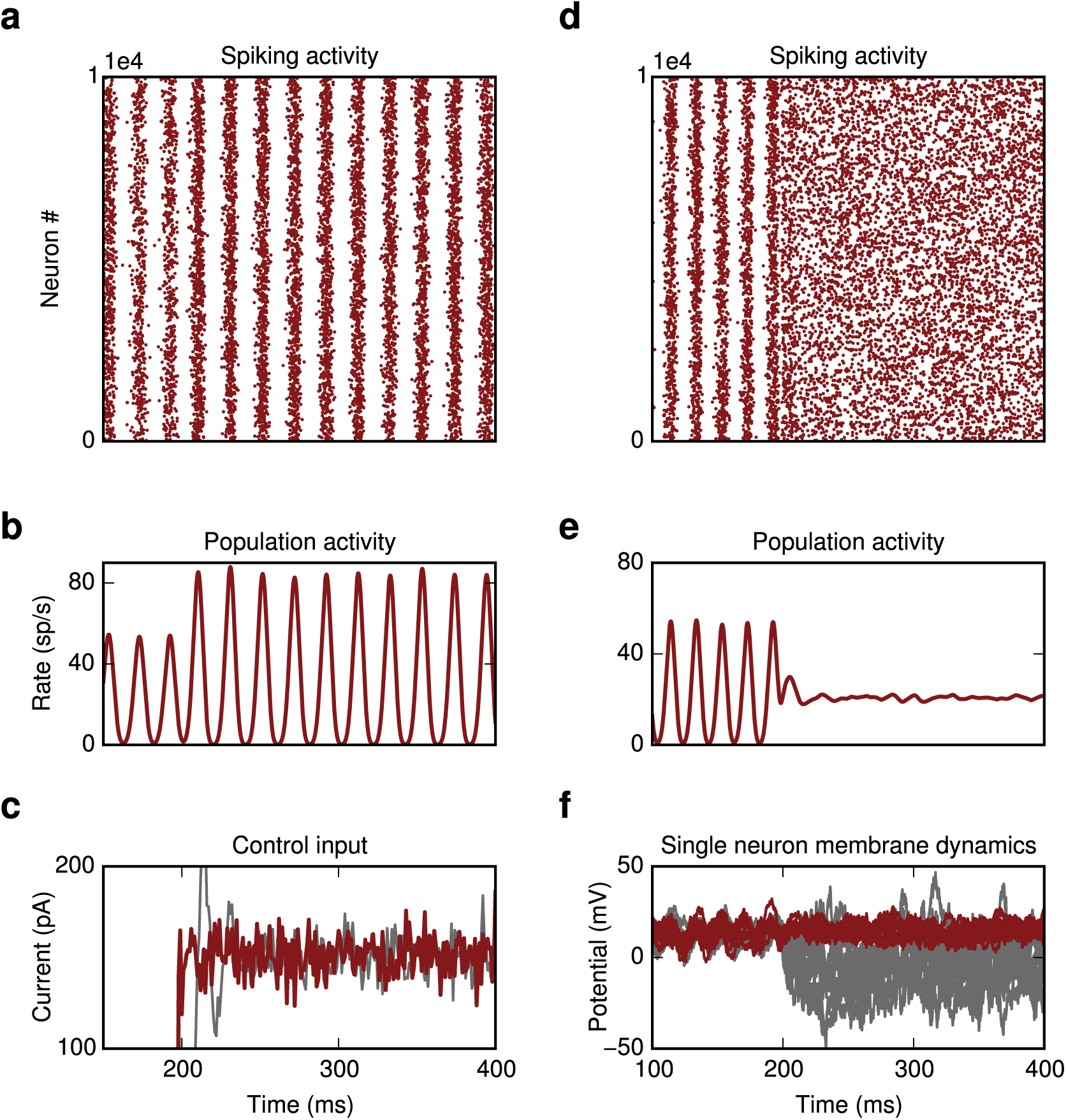
Noise injection (**A** and **B**) Injecting Gaussian noise with the same mean and variance as the control signal does not result in suppression but rather in an enhancement of oscillations. (**C**) Current injected into the somata of the neurons. Grey: Gaussian noise, red: DFC signal used in fig 2a-d. (**D** and **E**) Injecting strong Gaussian noise, *σ* =14 mV, yields to suppression of the oscillatory activity. (**F**) The subthreshold dynamics of ten randomly chosen neurons reveal that this strong external noise results in very large fluctuations in the membrane (grey). By contrast, the fluctuations under DFC are significantly smaller (red).

## Recovery of network function

The detrimental effect of strong external noise became even more apparent when we studied the response of the network to incoming stimuli. We examined two scenarios. First, we tested how a series of incoming pulse packets composed of randomly distributed spikes are processed by the SNN. We evaluated the network response by the area under the curve (AUC, see methods) for each of the following network states: AI, SI under DFC and SI under noise stimulation. A high AUC value reflects better separability of two conditions. It is evident that the AUC in the AI state and in the DFC condition is close to unity indicating that both conditions are comparable in terms of stimulus separability (**Figure 5A,B**). By contrast, when the SI oscillations were suppressed by the injection of strong external noise the AUC dropped significantly. That is, DFC, in contrast to strong noise stimulation, does not impair the ability of the network to detect incoming stimuli. These results suggest that processing of incoming signals either locally or by downstream areas is feasible in a DFC scheme.

The previous test only captured the *firing-rate* coded processing that the network may be performing. Therefore, next we tested how DFC affects *temporal* aspects of the network response. To this end, we provided external correlated inputs to all stimulated neurons and measured the spike train similarity in the network response. We computed the spike distance *D*^33^ that captures the time-resolved degree of synchrony between individual spike-trains (see methods). Again DFC did not impair the temporal processing as indicated by a clear separation of the two clusters during baseline *D_B_* and stimulation *D_S_* (**Figure 5C**). For external noise, however, the two distributions of values strongly overlapped, showing that aspects of temporal processing as measured by pairwise synchrony are clearly compromised when the SI state is disrupted by open-loop noise injection.

**Figure 5.**
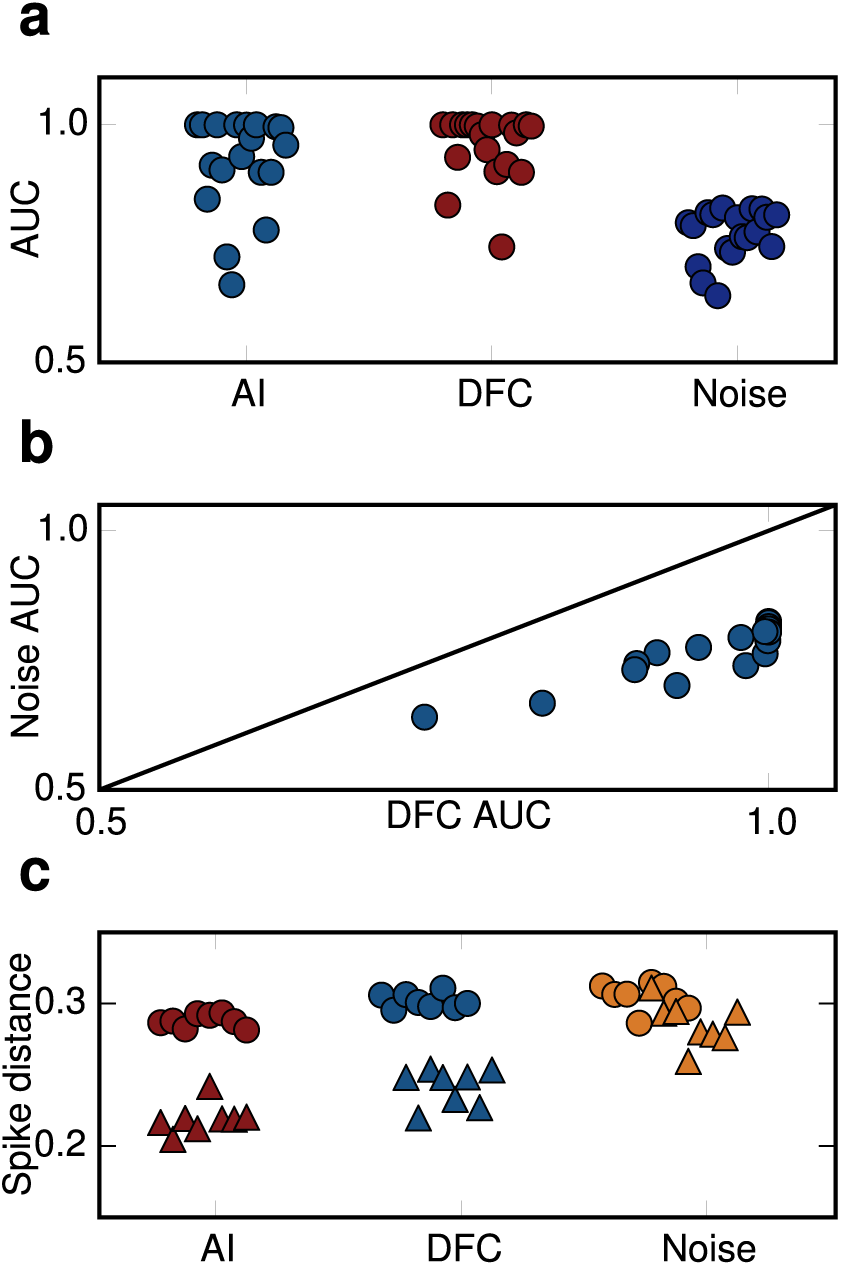
Recovery of rate and temporal based computations. (**A**) AUC values indicating how well the population rate response to incoming stimuli estimated for different time scales (different dots within each group) is separated from baseline activity for three different scenaria. (**B**) AUC values for DFC are systematically higher and close to one compared to noise injection. (**C**) A clear separation of spike distance values between baseline (dots) and response to incoming stimuli (triangles) indicates that temporal aspects of computations during DFC are comparable to the AI state. By contrast, noise injection leads to a strong overlap of the two distributions resulting in impaired temporal processing.

## Mechanism of DFC

The above two results clearly demonstrate that DFC has multiple advantages compared to the open-loop noisy stimulation. DFC does not only suppress SI activity steering the network to an AI regime, it also facilitates the recovery of the network’s ability to process stimulus related information. From its design it is evident that DFC effectively counteracts the increase in coupling strength, which is one of the main causes for the emergence of SI activity. Indeed, the goal of the DFC design was to move the poles of the system at, or close, to their original positions. Ideally, the stimulation kernel *M* would match the synaptic kernel *S* with *d_c_* = *d* and the amplitude of the control gain *K* would be tuned to match the pathological increase of the coupling strength Δ*J*. If this were the case, DFC would completely eliminate the effects on the mean recurrent input. This is evident if we consider the modulation to a perturbation in the average input to a neuron

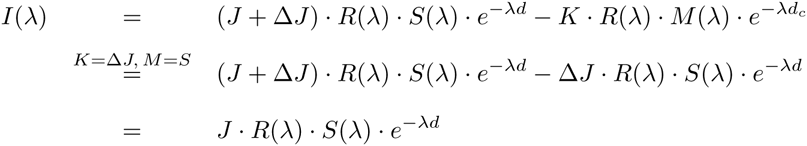

That is, under DFC the effects of Δ*J* are not visible in the perturbed current term. In practical applications a perfect match between the control parameters (*K,d_c_, M*) with the synaptic values is not feasible, because the exact shape of the synaptic kernels are not known a priori and have to be estimated. Nevertheless, within a certain reasonable range of parameters (see also section “stable control domains”), DFC still places the eigenvalues close to their initial position before the onset of pathology. Therefore, as we showed above, aspects of both rate and temporal coding that the network may be performing are recovered.

## Effects of neuronal and synaptic response function

The understanding of the exact mechanisms by which DFC suppressed SI activity allowed us to precisely investigate how the neuron and synapse response function *R* and *S* respectively influence the stability of the closed-loop system. To this end, we used again the mean-field approximation, because it incorporates explicit expressions for *R* and *S*. In general, the neuron response *R* depends on the specific neuron model as well as on the external input. Here, we did not change the neuron model, but altered the external Gaussian white noise input by using different values for the mean and variance (*μ, σ*^2^). We then assessed the stability of the system. It is apparent that for a given pair of coupling and control parameters (*J, d*) and (*K, d_c_*), respectively, the system becomes unstable as we move in the two dimensional parameter-space (**Figure 6A**). For meaningful comparison we used (*μ, σ*^2^)-combinations that yield constant rates. In the ideal case where *M*(*λ*) = *S*(*λ*) and *d_c_* = *d* equation (2) becomes:

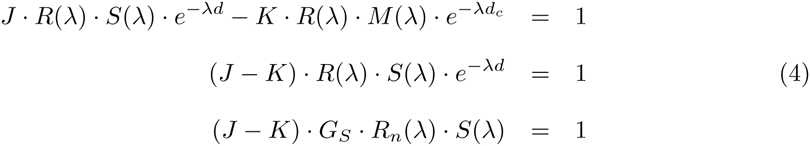

where *G_s_* is the slope of the ‘f-I curve’ at the operating point or static gain and *R_n_* the normalized neuron response (see methods). The critical effective coupling is then given by

**Figure 6.**
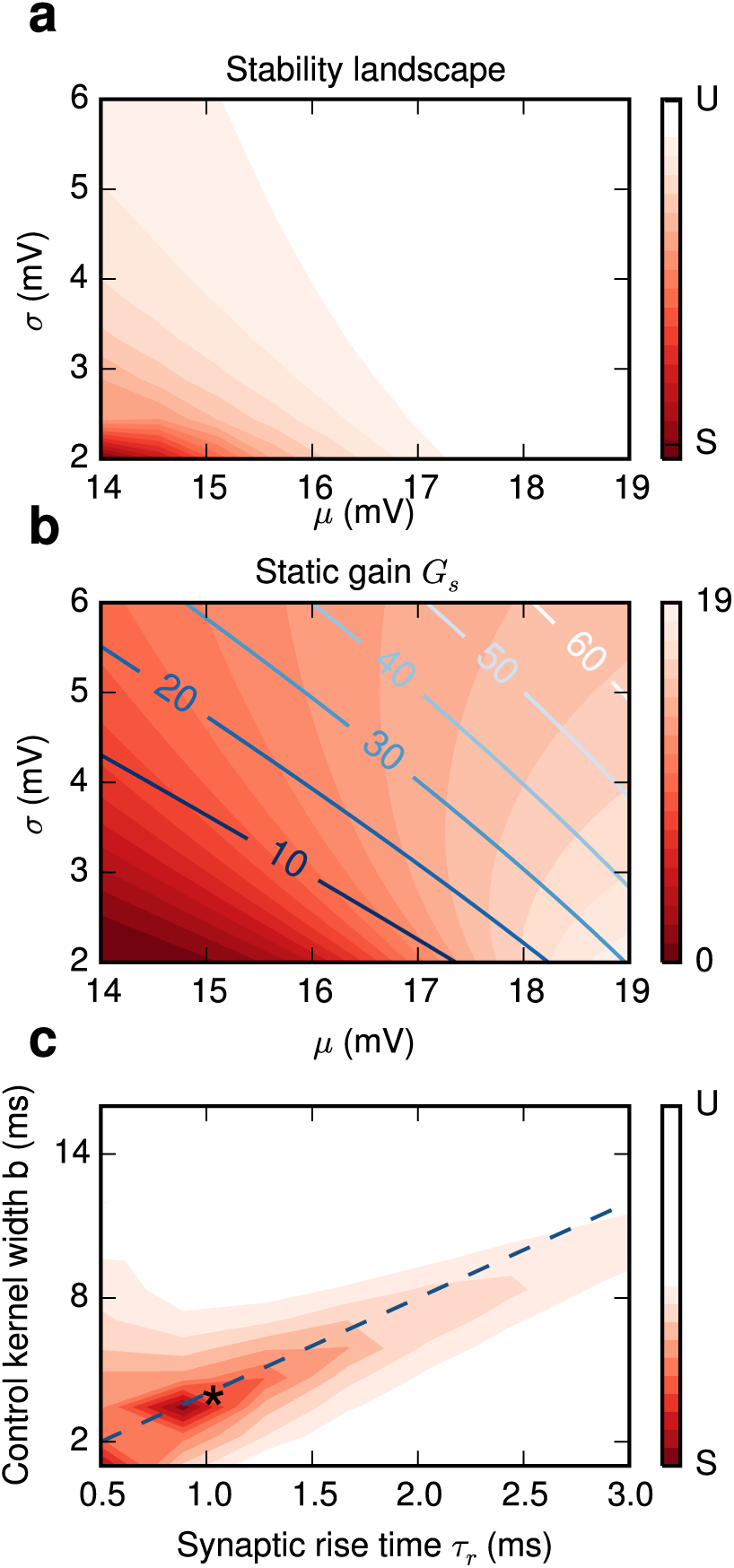
Effects of neuronal and synaptic response functions on stability. (**A**) For fixed network and control parameters (J,d) and (K,*d_c_*) respectively the stability of the closed-loop system changes with the operating point (*μ, σ*). (B) The static gain *G_s_* is the dominant factor for stability (see main text and Figure 6 - figure supplement 1). Lighter shades of red correspond to higher gain *G_s_.* The gain changes significantly even along the constant firing rate lines (blue lines, 10–60 sp/s). (**C**) The most stable control is achieved if the difference Δ*_d_* = (*d* + 2*τ_r_*) − (*d_c_* + *b*/2) between effective delays of control and synaptic kernels is minimized. The blue dashed line corresponds to *d* = *d_c_* where the optimal kernel width is *b* = 4*τ_r_*. It predicts correctly the stable regime for the range where *atan*(*ωτ_r_)* ≈ *ωτ_r_* (see Methods). In our simulations the effective control delay is *d_c_,_eff_* = 7 ms, which is very close to the optimal value (star). In panels (**A**) and (**C**) the stable and unstable regimes are marked by red and white colors respectively.

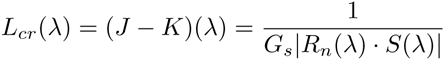

As we move along the constant output firing rate lines both *G_s_* and |*R_n_*(*λ*)| increase (**Figure 6 - figure supplement 1A,B**) leading to a decrease of *L_cr_*. The changes in *G_s_* are significantly larger than those in |*R_n_*(*λ*)|, implying that the static gain is the dominant factor that affects stability. The changes in *|S*(*λ*)| are negligible (**Figure 6 - figure supplement 1C**). This is expected, because the frequency range we are interested in is much smaller than the cut-off frequency of the synaptic filter *ω* < *ω*_3_*_db_.* Thus, when the system operates in a dynamic regime in which single neurons’ responses have a higher gain the control domains shrink and the range of *K* values that stabilizes the system decreases.

Next, we investigated the interaction between the synaptic *S*(*λ*) and the control kernel *M*(*λ*). The amplitude responses for different kernels do not vary significantly (**Figure 6 - figure supplement 2**) Therefore, the important factor that influences stability is the phase difference or, alternatively, the difference Δ*_d_* between the effective delays of the synaptic *d_eff_* and the coupling kernel *d_c_,_eff_*. An optimal result is achieved if this difference vanishes (see methods)

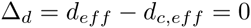

This point is illustrated for the case where *d_c_* = *d* + 1ms (**Figure 6C**). These results show that DFC does not depend strongly on the shape but rather on the effective delay of the kernel.

## DFC induced SI activity

Interestingly, the same control strategy can be used to induce or enhance rather than to suppress oscillations. Choosing appropriate control parameters to increase the effective coupling, i.e. selecting K to have the same sign as *J* (see methods), results in SI activity (**Figure 7**). This may be helpful for the treatment of symptoms in several pathological conditions that are characterized by impaired oscillations, e.g. gamma power decrease in schizophrenia^34^. Thus, DFC is a generic control approach that, depending on the particular situation, can be used both to quench or to enhance oscillatory activity.

**Figure 7.**
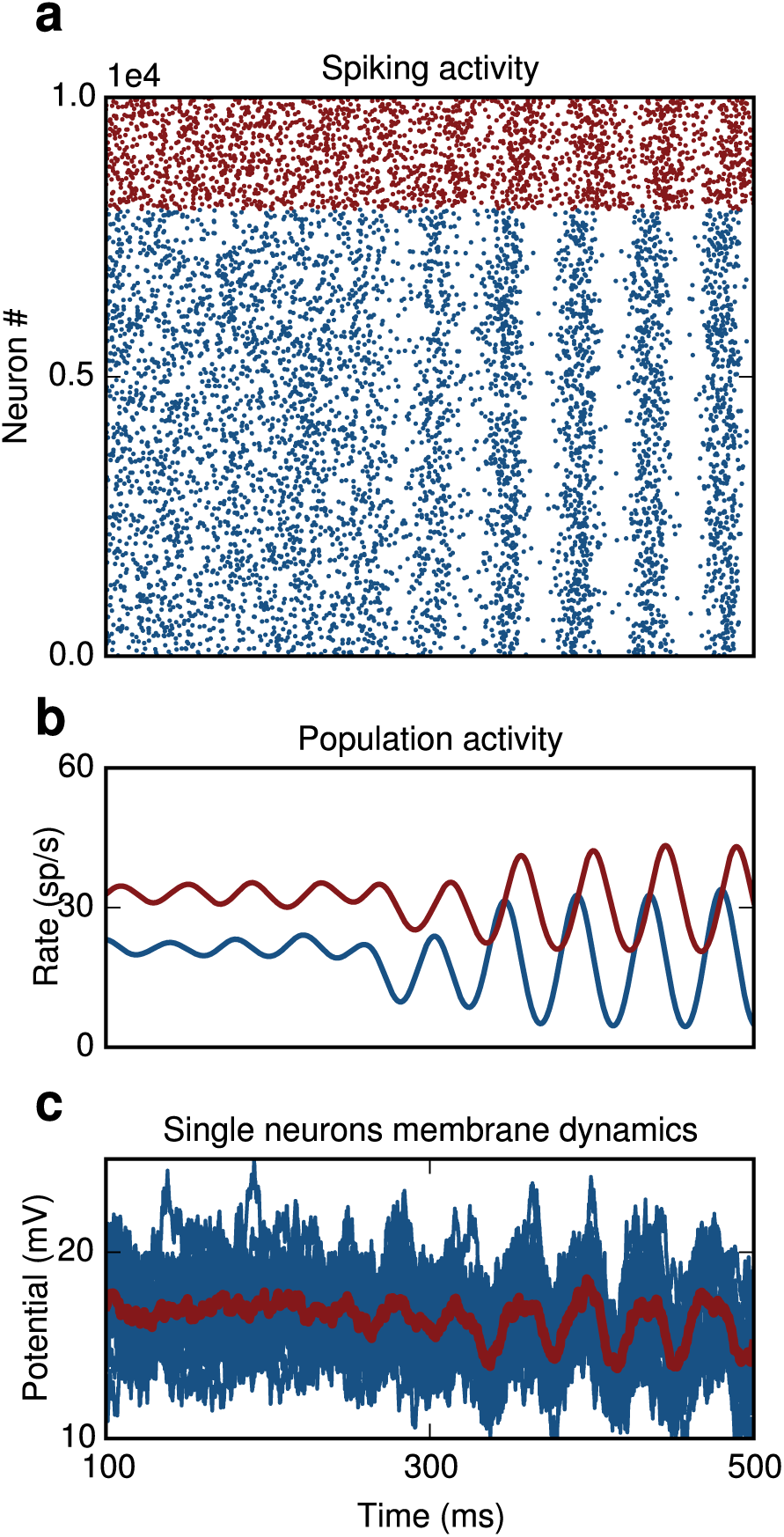
Creating oscillations. DFC can also be used to create rather than suppress oscillations. (**a**) E-I network. Switching on the controller at t=250 ms causes oscillatory activity. (**b**) Population activity of excitatory (blue) and inhibitory neurons (red). (**c**) Single membrane potential trajectories of ten randomly chosen excitatory neurons. Averaged trace of subthreshold dynamics is shown in red.

## Discussion

Open-loop stimulation has been the main non-pharmacological approach to control the symptoms in a wide range of pathological conditions. It has been successful in parts, but it often introduces clinical side-effects^35^. Moreover, it inherits the drawbacks from open-loop systems: (i) The stimulation profile is predetermined and is not adjusted to the clinically observed shortterm fluctuations in the patients’ symptoms^36^. In addition, stimulation is continuously applied even though it may not be always necessary (ii) The stimulation does not adapt to long-term changes of the system, e.g. structural alterations due to the progression of the disease. (iii) The operating point cannot be altered to deal with perturbations, caused, for instance, by a drift of the electrode lead^37^ (iv) External disturbances due to transient undesired signals are not being suppressed.

In contrast to this, closed-loop control can *by design* deal with all these situations. For this reason studies have started to investigate feedback-control both experimentally^2,21-23^ and theoretically^38^. The goal of the experimental work has been to demonstrate that closed-loop control is indeed effective, whereas the theoretical studies aimed at providing a deeper understanding of the underlying mechanisms.

Here, we provide a theory for DFC, a conceptually simple but powerful form of control,^5,6^ applied to the suppression of stochastic SI oscillations in SNN. These oscillations are generic, they occur in many brain areas and in multiple conditions^39^ and they emerge via a supercritical Hopf bifurcation^16^. Therefore the control objective was specific: to counteract this bifurcation. We provide a mean-field approximation to estimate the DFC parameters and confirm the analytical predictions in numerical simulations in purely inhibitory and in coupled excitatory-inhibitory SNNs.

We used two control approaches, direct and differential control, and demonstrated that both schemes are effective in suppressing oscillations. Consistent with previous findings^5,29^, our results reveal that differential control has two main advantages over direct control. First, the control domains are enlarged, which renders the selection of control parameters an easier task. Larger control domains implies increased robustness of the system both to perturbations in the parameters and to disturbances. This means that neither small deviations from the nominal values of 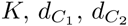 nor external signals compromise its stability. Second, in differential control the stimulation signal vanishes which translates to decreased power consumption. In clinical settings this is a highly desirable property and is, in fact, a basic requirement of any neuroprosthetic device.

The key advantage of the approach we presented here is that the system under control is being steered back towards its primary operating point (**Figure 8**). That is, DFC effectively decreases the synaptic coupling strength and, therefore, it counteracts the causes that originally led to the instability. This is obviously true only for the first-order statistics, because DFC does not counteract changes in the variance of the input that a random neuron in the network receives. Nevertheless, this is sufficient for the network to recover basic processing abilities both for rate and temporal coding schemes. Alternative approaches that rely on increased external noise are able to suppress oscillations^40^, but they do not allow the network to perform any meaningful computations. We think that a similar explanation is valid also for the traditional open-loop DBS. The exact mechanisms of this type of DBS are still debated^41^, but one of the reasons for the induced side-effects may be compromised information processing.

**Figure 8.**
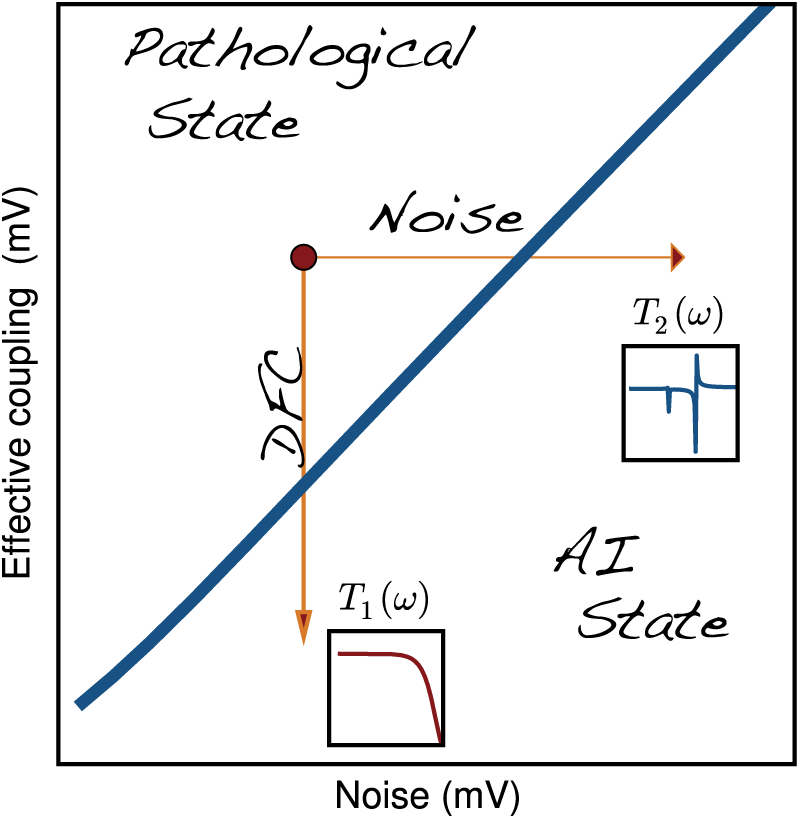
Recovery of network computations. DFC decreases the effective coupling between the neurons steering the system back to its original operating point and restoring the original transfer function *T*_1_(*ω*). Alternative approaches, e.g. noise injection may suppress oscillations, but they drive the system to a dynamical regime characterized by a different transfer function *T*_2_(*ω*) where physiological computations are impaired.

DFC suppresses oscillations in SNN not by decorrelating individual neurons, but rather by applying a common signal to all neurons that counteracts the mean input they receive from the network. Besides the many advantages described earlier, the utility of DFC lies in the fact that it is a very general control strategy, which does not depend qualitatively on lower level properties such as the specific coupling kernels of the connections. Neither does it depend qualitatively on the exact neuronal type (**Figure 2 - figure supplement 1**). Thus, DFC can in principle deal with more complex scenarios, e.g. heterogeneities in network and in neuronal properties. Nonetheless, the exact shapes of the neural and synaptic response functions do affect the system quantitatively and do modify the stability landscape. It is, thus, essential to have a good understanding of their precise contribution.

The results we presented provide a clear picture about how exactly the static gain of the neuron affects the size of the stable control domains. We also showed that the width rather than the shape of the control kernel affects the stability boundary. Further, we explained how the coupling strength and delay influence the overall stability landscape. These insights have besides the theoretical value a direct implication for the design of neuroprosthetic devices and are, therefore, of immediate practical and clinical relevance.

It is also important to address certain limitations in our approach. First, we assumed that we can stimulate neurons by current injection. In practice this is currently not possible, therefore incorporation of volume conduction models^42^ to describe the effects of external stimulation on individual neurons would be required. Second, we used the population average of single neuron firing as our state variable. Again, in more realistic settings an indirect measure of population activity such as a local-field potential (LFP) signal^2^ has to be used. Third, our theoretical analysis was based on a mean-field approximation that ignores fluctuations in the input. Analytical and numerical results were largely in a good agreement, but additional work is necessary to specifically deal with the fluctuations in the activity. Last, we had access to the relevant parameters required for the tuning of the controller. In real applications these parameters have to be estimated online from the recorded activity. The delay could be inferred from the frequency of the oscillatory activity. Inference of the coupling strength is less straightforward, but may still be feasible. Alternatively, once the delay is estimated, methods of adaptive tuning could be used to retrieve also the optimal control gain (**Supplementary figure 1**). Tuning the controller is in general a difficult problem, even for open-loop DBS, and additional research in this direction is required.

## Differences from previous work

View studies have addressed the problem of suppressing oscillations in neural activity (see^38^ for a detailed review). They are based (i) on population dynamics^43^^15^ (ii) on detailed single neuron descriptions^13^ or on combinations thereof^14^. These approaches have their merits, but they come with limitations: (i) the parameters cannot be directly mapped to experimental measurable quantities (ii) it is not clear if the results scale to large network of neurons.

The approach that we presented here is a trade-off between biophysical realism and analytical tractability. We used the LIF model, which captures single-neuron dynamics to a sufficient degree, while at the same time allows computationally efficient simulations of large networks. We applied DFC that was originally proposed in the context of chaotic systems as a method to control unstable periodic orbits^5^. It was later used to control coherence^44^ and to suppress synchronous activity in networks in which the neurons themselves act as oscillators^10-12^.

Here, we did not use simplified population dynamics or phase oscillators. Instead, we used spiking neurons that fire irregularly and are nevertheless able to generate oscillations. We also used realistic models of synaptic dynamics and were therefore able to explicitly study their contribution to stability. This allowed us to design an appropriate control kernel, which resulted in increased control domains. In addition, by using a mean-field theory that explicitly incorporates the synaptic and neuronal response functions we could study their contribution in a systematic way. The neuronal response function enabled us to investigate the influence of external and recurrent inputs and to relate them to experimentally measurable quantities. Indeed, as we showed above, the statistics of the mean field for activity states with very similar firing rate profiles may be significantly different affecting stability. Therefore, feasible measurements of the population activity can be directly used to characterize the operating point of the network and to fine-tune the control parameters to achieve the desired results.

## Conclusions

We used DFC, a relatively simple form of control that includes only a proportional gain term, because it is still possible to analytically study the stability of the closed-loop control system. More sophisticated control strategies could further increase the performance of the system. They come, however, at the price of increasing the number of control parameters that have to be estimated and of increasing complexity precluding a formal proof of stability. The approach we presented here spans multiple levels of analysis of neuronal dynamics, enabling an understanding of how the control stimulus interacts with both low-level synaptic and high-level properties of the population activity to influence stability. At the same time the complexity of the controller is kept low to be of practical relevance. Thus, here we have provided a general conceptual framework for future studies that address both theoretical and practical aspects of closed-loop control in neuronal systems.

## Methods

### Numerical simulations

We use networks of *N* LIF neurons randomly connected with a probability of *ε* = 0.1. Thus each neurons receives *C* = *εN* connections from other neurons in the network. For the purely inhibitory network we use *N* = *N_I_* and for the coupled excitatory-inhibitory case *N* = *N_E_* + *N_I_.* The subthreshold dynamics of a neuron *i* in the network is given by

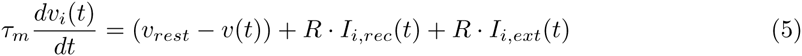

where *τ_m_* is the membrane time constant and *v_rest_* is the resting potential. The recurrent input term

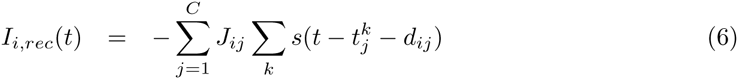

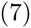

describes the total synaptic current arriving at the soma due to presynaptic spikes. Each presynaptic spike causes a stereotypical postsynaptic current *s*(*t*) modeled as an *α*-function^45^

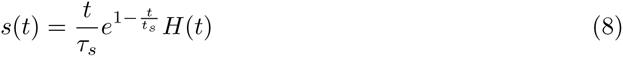

where *τ_s_* is the synaptic time constant and *H*(*t*) the Heaviside function.

The double sum in equation 5 runs over all firing times 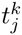 of all presynaptic neurons 1, 2,…, *C* connected to neuron *i*. For all connections in the network we use the same synaptic coupling strength *J_ij_* = *J/N* and the same connection delay *d_ij_* = *d.* The external input

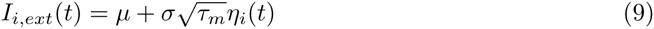

contains a mean term *μ* and a fluctuating term resulting from the Gaussian white noise *η_i_*(*t*) that is uncorrelated from neuron to neuron with < *η_i_*(*t*) >= 0 and < *η_i_*(*t*)*η_i_*(*t′*) >= *δ*(*t* − *t*′).

## Asynchronous state

In the stable asynchronous state the population activity is constant *v*(*t*) = *v*_0_. The mean recurrent input that each neuron receives is therefore also constant and given by

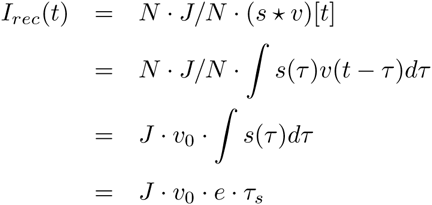

We study the stability of the asynchronous state following a linear perturbation approach^18^. A small oscillatory modulation of the stationary firing rate *r*(*t*) = *r*_0_ + *r*_1_*e*^−^*^λt^* with *v*_1_ ≪ 1 and *λ* = *x* + *jω* where *ω* is the modulation frequency leads to corresponding oscillation of the synaptic current

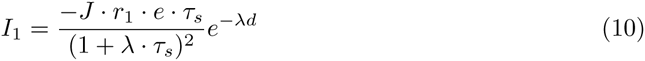

The firing rate in response to an oscillatory input is given by

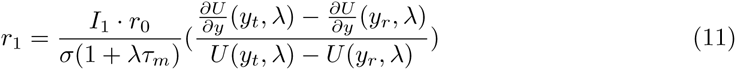

The function *U* is given in terms of combinations of hypergeometric functions

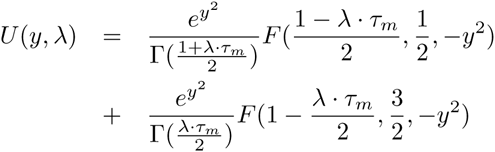

In a recurrent network the modulation of the firing rate and the modulation of the synaptic input must be consistent. Combining (10) and (11) we get

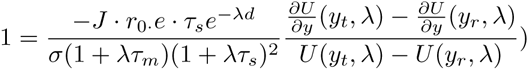

which we write as

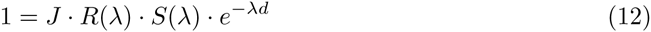

where the terms

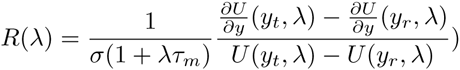

and

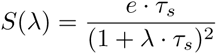

describe the neuronal and synaptic response functions respectively. The negative sign of *J* is absorbed in the phase of *S*(*λ*).

The critical coupling values at which modes have marginal stability with frequency *ω_i_* can then simply be computed by

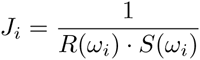

The smallest value *J_cr_* = *min*{*J_i_*} is the critical coupling at which the first complex pair of eigenvalues crosses the imaginary axis and the system becomes unstable. In the case of the inhibitory network for *m* = 14 mV and *σ* = 6 mV we have *J_cr_* = 115 mV. In the simulations we used for the coupling between two neurons *i* and *j*, *J_ij_* = 0.2mV thus the total coupling is *J* = *C · J_ij_* − 1000 · 0.2 mV= 200 mV *>J_cr_* (**Figure 2A-D**).

### Stability

The eigenvalues *λ* of the self-consistency equation (12) determine the stability of the system. If for all solutions the real part is negative, *Re*{*λ*} < 0, then the system is stable otherwise it is unstable. The stability border, *λ* = *jω*, is characterized by the occurrence of a supercritical Hopf bifurcation. At this point the population activity will be oscillatory with frequency *ω*.

## Delayed feedback control

In the simulations we implement DFC by recording and stimulating all neurons in the network. The subthreshold dynamics of a neuron *i* with DFC is given by

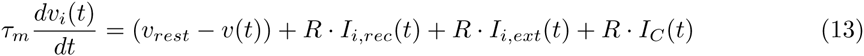

where *I_C_*(*t*) is the control input. Note that *I_C_*(*t*) is identical for all neurons in the network given by

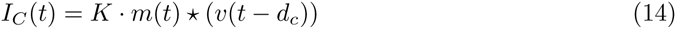

where *v*(*t*) is the instantaneous population activity at time *t* and * denotes the convolution operation 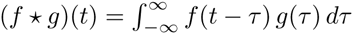. We used as control kernel *m*(*t*) a box function

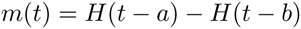

where *H*(*t*) is the Heaviside function

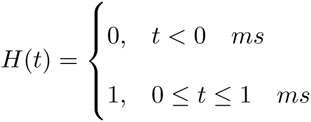

Thus the control input *I_C_*(*t*) was updated in steps of 1ms.

### Direct control

In the case of direct DFC a modification of the self-consistency equation (12) yields

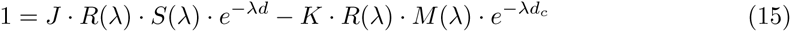

where *M*(*λ*) describes the control kernel in the frequency domain and the negative sign captures the fact that the control stimulus counteracts the effects of the synaptic coupling *J*.

The box function has response characteristics given by

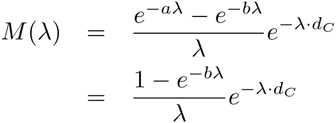

where *a* = 0, *b* is the width of the kernel and *d_c_* is the control delay.

### Differential control

In differential DFC we the control input *I_C_*(*t*) is a function of the difference between two time-delayed versions of the population activity *v*(*t*). It is given by

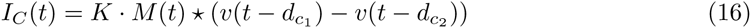

In this case the self-consistency equation (12) is modified to give

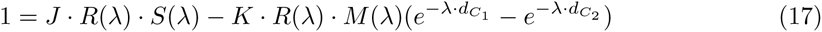

### Rate compensation

In all simulations we adjust the mean *μ* and variance *σ* of the external input to the neurons such that the firing rates are approximately equal for all conditions, that is

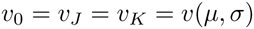

where *v*_0_, *v_J_*, *v_K_* are the firing rates of the uncoupled, coupled and network under DFC respectively. In this way we can exclude any effects due to changes in the firing rates.

### Stability analysis

The eigenvalues *λ* of the self-consistency equations (15) and (17) determine the stability of the system. We compute for both direct and differential control the real part of the rightmost eigenvalue *Re*{*λ*_1_} that determines stability. We use the (*I_K_, d_c_*)-parameter pair with *I_K_* ∈ [0,300] mV and *d_c_* ∈ [0,30] ms. The second delay term in differential control was in both cases *d_c_*_2_ = 1 ms.

### DFC induced SI activity

If the control gain *K* has the same sign as the synaptic coupling *J* and the control delay is chosen to be close to the synaptic delay, *d_c_* ≃ *d*, then the effective coupling in the network increases resulting in SI activity (**Figure 7**). In this case the self-consistency equation is given by

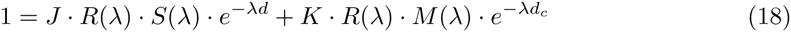

### Static gain

In the frequency domain the neuron response function *R*(*f*) is simply the Fourier Transform of the impulse response *h*(*t*), i.e. 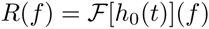. The impulse response *h*(*t*) can be separated in two parts

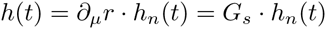

where *h_n_*(*t*) is the normalized impulse response such that 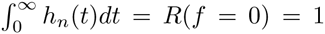 and *G_s_* = *∂_μ_r* is a constant term that we denote as the static gain of the response. It corresponds to the slope of the ‘f-I’ curve at the operating point and captures the ‘susceptibility’ of the rate to small changes in the mean *μ.*

We can rewrite the self-consistency equation 4 as

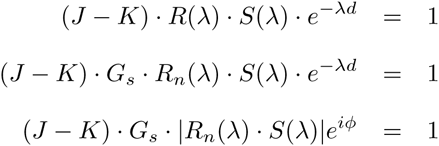

where we have separated the complex expression *R_n_*(*λ*) · *S*(*λ*) in an amplitude and phase part. Splitting this complex equation in two real ones we get

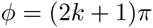

and

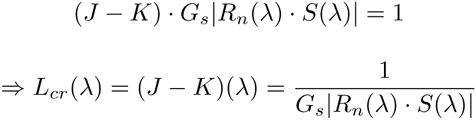

The first equation gives us the critical frequencies *ω_cr_* for which the modes exhibit marginal stability, i.e. *λ* = *iω_cr_*. From the second equation we can compute the corresponding effective critical coupling *L_cr_*(*ω_cr_*).

### Control kernel dependence

To study how stability depends on the control kernel we write again the self-consistency equation

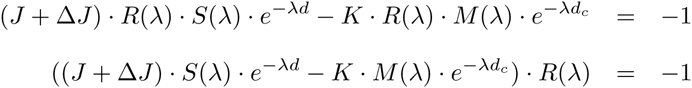

where Δ*J* is a pathological increase in the synaptic coupling that the controller needs to counteract. Separating amplitude and phase we get

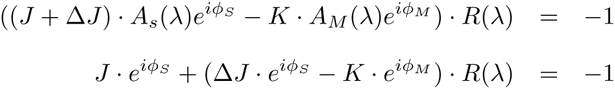

where we used *A_S_*(*iω*)) = *A_M_*(*iω*) = 1, which is valid for the frequency range that we are interested in *ω* < 300 rad (**Figure 6 - figure supplement 2**). To counteract the increase Δ*J* we need to minimize the effective coupling term

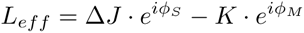

which leads to

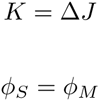

For the control kernel we use a box function defined as

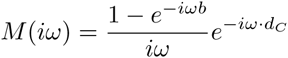

with phase

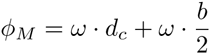

whereas the synaptic kernel is the *α*-function

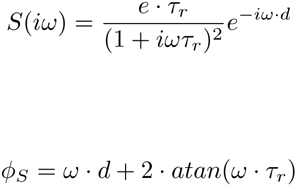

Thus

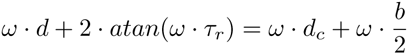

and if we assume that *atan*(*ω · τ_r_*) ≈ *w · τ_r_* then

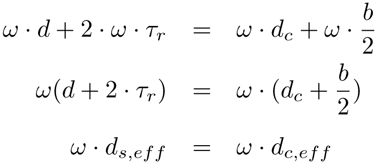

Thus, the most optimal control is achieved when the effective delays of synaptic and control kernel *d_eff_* and *d_c_,_eff_* respectively are identical

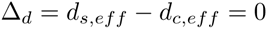

In our simulations we used *d* = 5 ms and *τ_r_* = 1ms, thus *d_s_,_eff_* = 7ms. For the controller we used *d_c_* = 6.5 ms and *b* = 1ms, thus *d_c_,_eff_* = 7ms.

## Recovery of network function

### Response to random pulse packets

We computed the response of the network to incoming stimuli that arrived in form of random Gaussian pulse packets. A pulse packet was composed of a predefined number of spikes *n_pp_* = 100 with normally distributed random displacements *t_i_* ~ *N* (*μ* = 0, *σ_pp_* = 10) from the center time *t_c_* of the pulse. It was fully defined by the tuple (*t_c_, n_pp_, σ_pp_*). In total, ten pulse-packets with center times *t_i_* = (200*i* + 200) ms and 1 ≤ *i* ≤ 10 were applied to 100 randomly chosen neurons in an inhibitory network of size *N* = 1000. We computed the population response at time points *t_i_* + 10 ms and compared it with the population activity during baseline at time points *t_i_* + 100 ms. For this, we performed a receiver operating characteristic (ROC) analysis evaluating the true positive and false positive rate for various thresholds. We then computed the area under the ROC curve (AUC), which indicates how well the response can be distinguished from baseline activity. An AUC value of 1 means full separability of the two activity states, whereas an AUC value of 0.5 indicates full overlap of the activity sampled during the two different conditions. We computed the AUC values for three different scenarios: (i) physiological AI state (ii) pathological (oscillatory) state controlled with DFC (iii) pathological (oscillatory) state with noise. The results are shown in **Figure 4A,B**.

### Response to common input

We defined a spike train of *n_ST_* = 500 equally spaced spikes in a window of *T_ST_* = 50 ms. Ten copies of exactly the same spike train with time onset *t_i_* = (200*i* + 200) was provided as input to 100 randomly chosen neurons in an inhibitory network of size *N* = 1000. Thus, in this scenario all stimulated neuron received *identical* input during the stimulation periods. However, in this case we were interested in the temporal aspects of the network response. To this end, we measured the synchrony between the spike trains of all neurons in the network using the SPIKE-distance metric^33^. The SPIKE-distance is a measure of (dis)-similarity which allows for a time-resolved analysis and can track instantaneous changes. We computed the multivariate SPIKE-distance *S* both during the 50ms of stimulation (*S_ST_*) and also during 50 ms of baseline activity (*S_BL_*). We then computed the temporal average for the stimulation

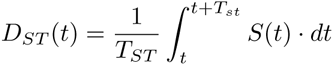

and for the baseline *D_BL_*(*t*) = *D_ST_*(*t* + 100). The results, again for the three scenarios described above (AI, DFC, noise) can be seen in **Figure 4C**.

### Oscillation index

We estimated the discrete power spectral density *P*(*ω*) of the population activity *r*(*t*) using the standard Fast Fourier Transform (FFT) method. We then computed the total power in the range [0, 250/*π*] rad

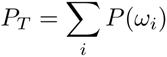

and used *log*_10_*P_T_* as a descriptor of oscillation strength.

## Author Contributions

I.V. and A.K. conceived the study. I.V. performed the analytical and numerical computations. T.D. assisted with the analytical computations. A.K., T.D. and A.A. participated in discussions. I.V and A.K. wrote the manuscript.

## Competing Financial Interests

The authors declare no competing financial interests.

Figure supplement 1. DFC control in heterogeneous networks.

Figure supplement 1. Static gain.

Figure supplement 2. Control kernels.

## Supplementary Information

### Effects of neural heterogeneity: bursting neurons

We test how well DFC performs under neural heterogeneity by substituting a fraction of regular spiking neurons with bursting ones. This particular choice is motivated by the fact that around 30% of neurons in the GPI/STN (,^46^H. Bergman: personal communication) exhibit bursting activity. We used a novel implementation of a bursting neuron type, in which the f-I curve of the neuron is not affected by the generation of bursting spike patterns (Saharasnamam, Vlachos, Aertsen, Kumar, under revision). It is evident that the existence of bursting neurons does not decrease the efficacy of the controller, which upon activation sufficiently suppresses oscillations (**Figure 2 - figure supplement 1**). These results suggest that DFC is effective in a wide range of networks where single neuron properties may be of secondary importance. Indeed theoretical work on the control of time-delayed system suggests that DFC is applicable to any system that undergoes a supercritical Hopf bifurcation as well as other types of bifurcations.^6,47^ Thus, for the application of DFC on the mean-field of the network activity neuronal heterogeneity does not pose a serious problem, which renders DFC particularly relevant for neuroprosthetic applications.

**Figure 2 - figure supplement 1.**
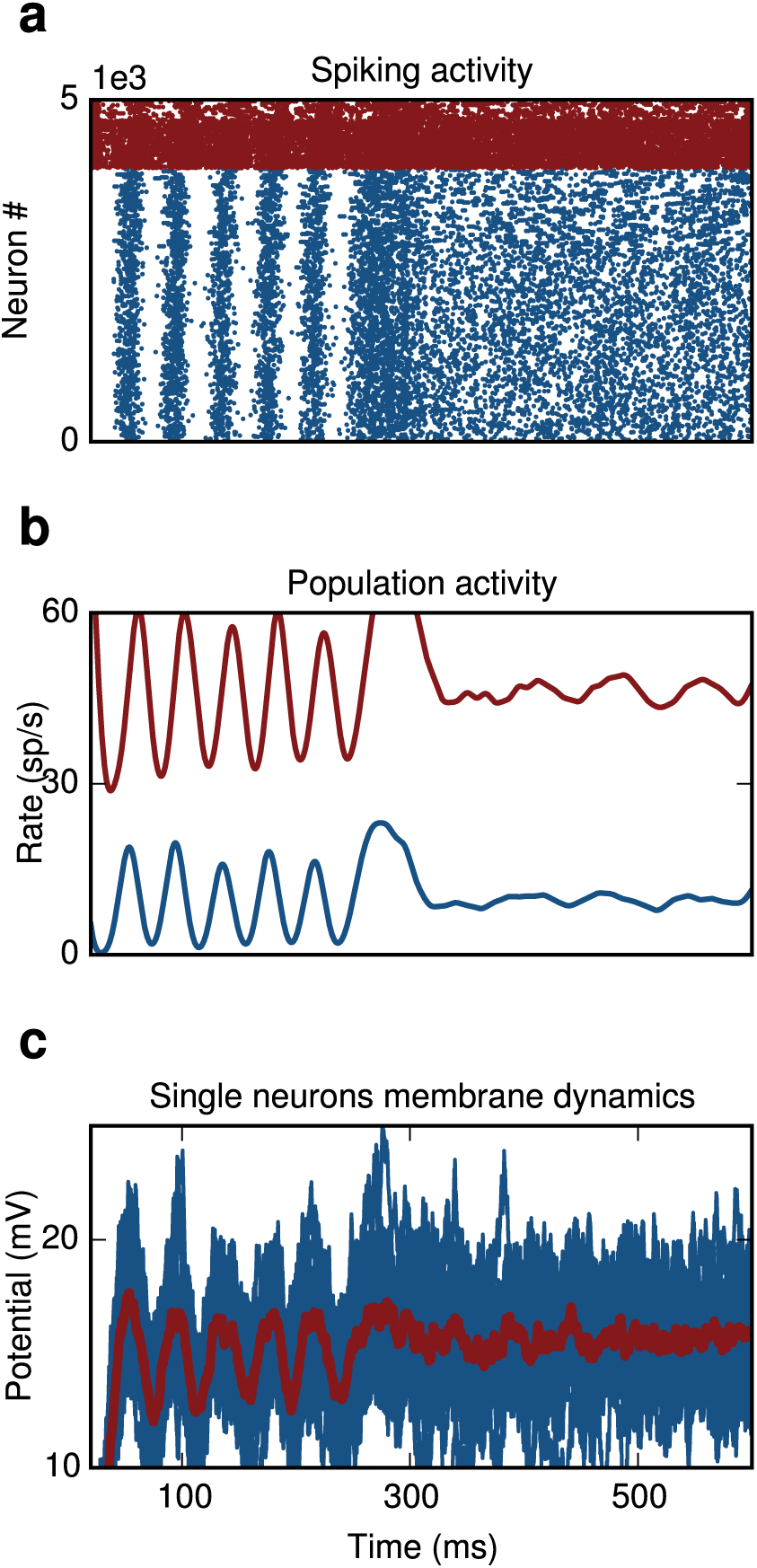
DFC control in heterogeneous networks. (**a**) Raster plot. Replacing regular spiking by bursting neurons (top 30% in excitatory and inhibitory population) does not compromise the effects of control. (**b**) Population activity of E-neurons (blue) and I-neurons (red). (**c**) Single membrane potential trajectories of ten randomly chosen E-neurons in the network. The averaged trace of the subthreshold dynamics is shown in red.

**Figure 6 - figure supplement 1.**
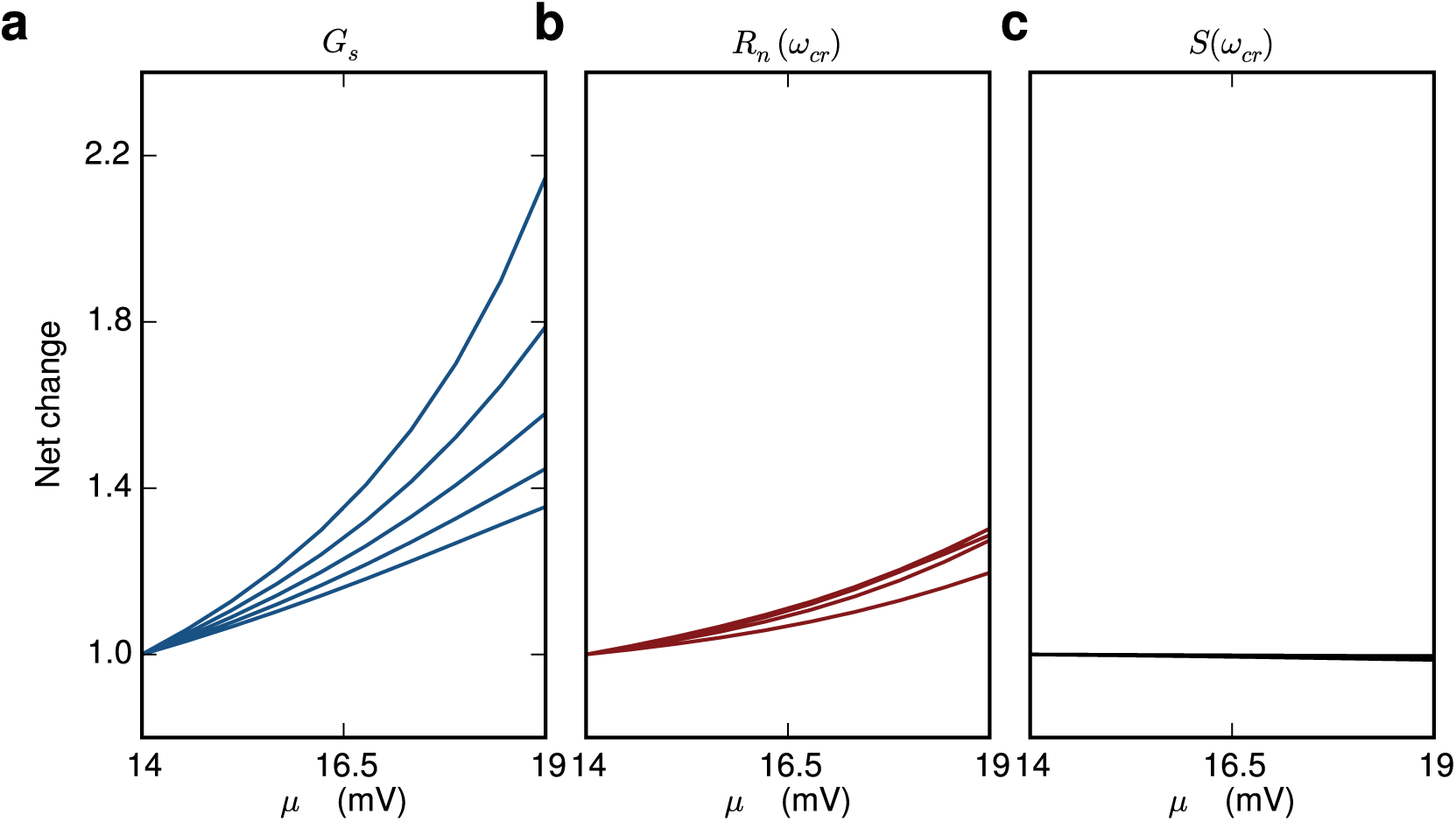
Static gain. In order to maintain constant firing rates in each population we increase the mean input *μ* while decreasing the variance of the input *σ*. (**a**) Moving in the state-space while maintaining the firing-rates yields significant changes in the static gain *G_s_*. (**b**) The changes in the normalized neuronal response *R_n_* are modest (**b**) and the changes in the normalized synaptic response *S_n_* are negligible (**c**).

**Figure 6 - figure supplement 2.**
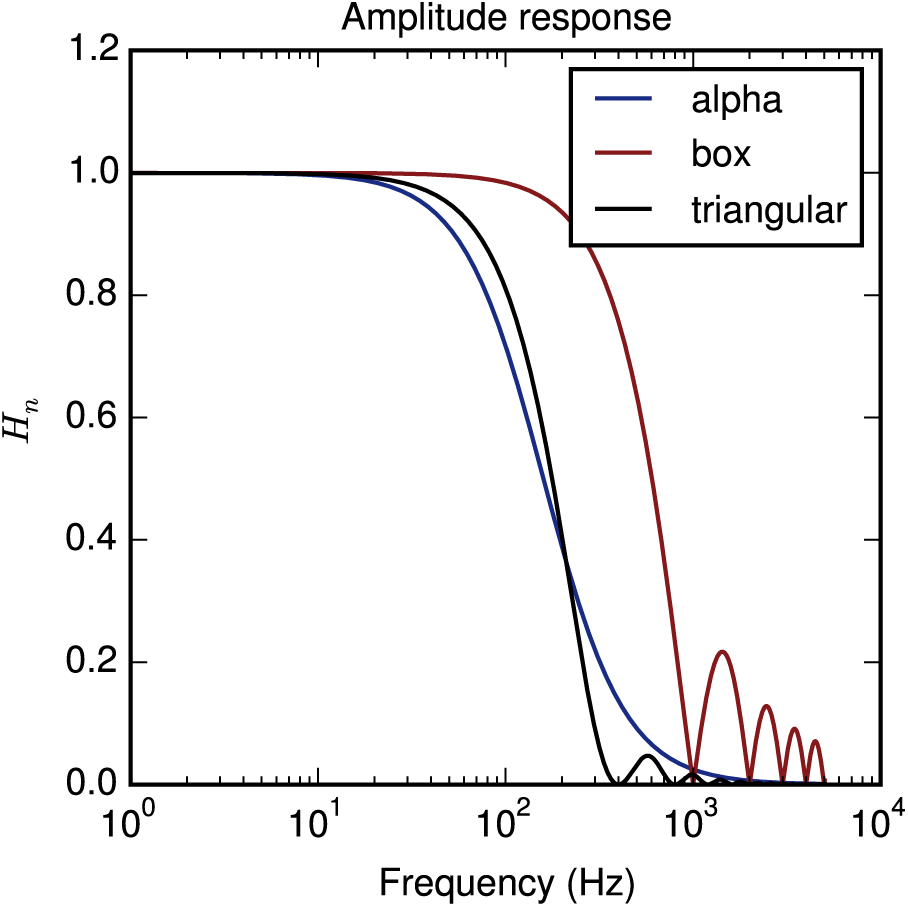
Control kernels. **S**_n_(*ω*) Various control kernels have very similar amplitude responses for the relevant frequency range f<100 Hz.

**Supplementary figure 1.**
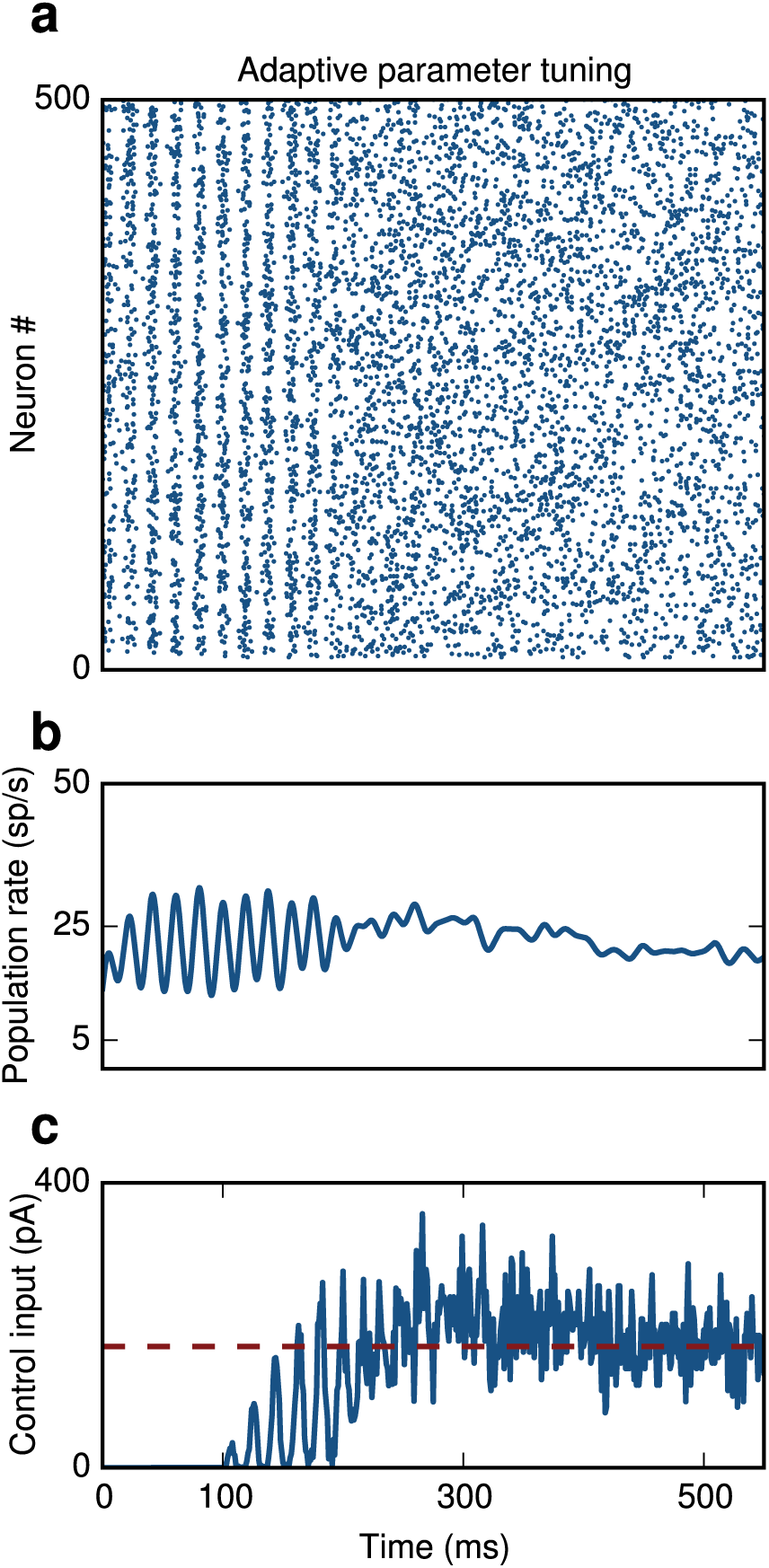
Adaptive parameter tuning. An adaptive procedure is used to find the optimal value for the control gain K. (**a**) I network. Switching on the controller at t=100 ms results in suppresion of oscillations. (**b**) Population activity of inhibitory neurons (blue). (**c**) The algorithm converges to an optimal value (red dashed line) for the control input within 200 ms after initiation of the procedure.

**Supplementary Table I:** Model parameters

| **Neuron Parameters** |  |  |
| --- | --- | --- |
| Spike threshold | $u_{th}$ | 20 mV |
| Reset potential | $u_r$ | 14 mV |
| Resting potential | $u_{rest}$ | 0 mV |
| Refractory period | $t_{ref}$ | 1 ms |
| Membrane capacitance | $C_m$ | 250 pF |
| Membrane time constant | $\tau_m$ | 10 ms |
| **Synaptic Parameters** |  |  |
| Synaptic rise time | $\tau_r$ | 1 ms |
| Synaptic decay time | $\tau_d$ | 1 ms |
| Synaptic weight | $J$ | |
| Synaptic delay | $d$ | 2 ms |
| **Input Parameters** |  |  |
| <u>Inhibitory network</u> |  |  |
| Input, mean - I population | $\mu_I$ | 14 mV |
| Input, variance - I population | $\sigma_I$ | 6 mV |
| <u>Excitatory-Inhibitory network</u> |  |  |
| Input, mean - E population | $\mu_E$ | 20.8 mV |
| Input, mean - I population | $\mu_I$ | 16.5 mV |
| Input, variance - E population | $\sigma_E$ | 20.8 mV |
| Input, variance - I population | $\sigma_I$ | 16.5 mV |
| **Network Parameters** |  |  |
| <u>Inhibitory network</u> |  |  |
| Number of inhibitory neurons | $N_I$ | 10,000 |
| <u>Excitatory-Inhibitory network</u> |  |  |
| Number of excitatory neurons | $N_E$ | 8,000 |
| Number of inhibitory neurons | $N_I$ | 2,000 |
| **Control Parameters** |  |  |
| <u>Inhibitory network</u> |  |  |

